# Application of the CPER reverse genetics system for genetic engineering of rabies virus

**DOI:** 10.1101/2025.10.22.683883

**Authors:** Yukari Itakura, Nijiho Kawaguchi, Koshiro Tabata, Gabriel Gonzales, Kei Konishi, Aiko Ohnuma, Itsuko Furuta, Naoto Ito, Shinji Saito, William W. Hall, Yasuko Orba, Hirofumi Sawa, Michihito Sasaki

## Abstract

Reverse genetics (RG) systems are essential tools for basic virological studies and applied studies using engineered recombinant viruses in various research fields. While the circular polymerase extension reaction (CPER) has been widely applied to prepare a full-length infectious complementary DNA (cDNA) of positive-sense RNA viruses, its use for negative-sense RNA viruses (mononegaviruses) remains limited. Here, we report the first CPER-based RG system for rabies virus (RABV), a member of mononegaviruses. Infectious RABV was successfully rescued from cells transfected with helper plasmids and the CPER product, the assembled overlapping DNA fragments encoding the full-length viral genome cDNA and regulatory elements. Using this system, we generated wild-type, point-mutant, reporter-expressing, and chimeric RABVs, all of which retained their expected biological properties. Deep sequencing revealed that CPER-derived viruses occasionally harbor low-frequency mutations undetectable by Sanger sequencing, highlighting PCR-related artifacts as a limitation. In addition, CPER products with a pUC19 backbone could be directly applied for *E. coli* transformation and cloning of RABV full genome cDNA plasmids, offering a flexible, ligase-free cloning strategy for conventional RG. Our work establishes CPER as a versatile platform for engineering recombinant RABVs, facilitating rapid generation and genetic manipulation of RABV with potential applications for research on other mononegaviruses.

## Introduction

Reverse genetic (RG) systems have provided great advances in virology research by enabling the generation of recombinant viruses with desired gene modifications, including nucleotide mutations, reporter gene insertions, and genome fragment exchanges between two viruses to produce chimeric viruses. These systems have become indispensable for molecular analysis of viral biological characteristics, antiviral screening, live-cell imaging, and viral vector development^1–4^.

For positive-sense single-stranded RNA (ssRNA) viruses, the process of RG is initiated by transfecting optimal cells with either viral genomic RNA or a plasmid encoding the full-length viral cDNA. The host cell machinery then induces the translation of viral proteins and subsequent virus replication^5^. In contrast, negative-sense ssRNA viruses require genome replication and mRNA transcription by the viral proteins on the ribonucleoprotein complex provided by the virions^6,7^. Therefore, in the RG of negative-sense RNA viruses, exogenous viral proteins must be provided via helper plasmids to initiate viral genome replication and mRNA transcription^8–10^.

Plasmid-based RG for rabies virus (RABV), a mononegavirus, is a well-established and widely used approach, which is mediated by transfection of a full-genome cDNA plasmid and helper plasmids^8^. RABV RG techniques have made great contributions to elucidating viral factors critical for RABV replication mechanisms and virulence, as well as investigating virus replication dynamics^11,12^. This has also enabled the engineering of RABV live vaccines exhibiting specific phenotypes^13,14^. Taking advantages of its high neurotropic characteristics, RABV reporter vectors have played significant roles in neuroscience enabling visualization of neuronal circuits^15^. However, the construction of a full-length viral genome expression plasmid is often time– and cost-consuming. Furthermore, gene cloning techniques such as restriction digestion, ligation, and homologous recombination are sequence-dependent and sometimes necessitate artificial sequence modifications.

The circular polymerase extension reaction (CPER) method, first reported for flaviviruses, offers a plasmid-free RG alternative^16^. In CPER, overlapping DNA fragments encoding a full-length viral genome cDNA and regulatory elements such as promoters and ribozymes are assembled into a circular DNA molecule in a single PCR. Direct transfection of this CPER product into appropriate cells enables recovery of recombinant viruses. This system has been applied to various positive-sense ssRNA viruses, including flaviviruses, alphaviruses, caliciviruses, and coronaviruses, and has made significant contributions to research on SARS-CoV-2 during the COVID-19 pandemic^17–19^. Since then, CPER design and virus recovery conditions have been optimized to improve the CPER-based RG system for positive-sense ssRNA viruses by expanding the target virus species and increasing virus recovery rates^19–22^. Due to the characteristics of the CPER method, it is thought that this method is advantageous when working with non-segmented viral genomes; however, its usage with negative-sense ssRNA viruses, i.e. mononegaviruses, had not been reported until the recent recovery of a wild-type respiratory syncytial virus using CPER^23^.

In this study, we report the first CPER-based RG system for the mononegavirus RABV. This platform enables the rapid and flexible generation of recombinant RABV including wild-type, mutants, reporter expressing, and chimeric viruses and underscores the potential of CPER as a versatile method for mononegavirus engineering.

## Results

### Establishment of a CPER-based RG system for RABV

We designed five overlapping DNA fragments encompassing the entire genomic cDNA of RABV CVS strain, along with a linker fragment of linearized pUC19 vector coding a T7 promoter and hepatitis delta virus-derived ribozyme. These fragments were prepared by PCR using a RABV CVS strain cDNA plasmid (pCVS)^24^ as a PCR template with specific primers (**Fig. 1A, 1B, and Table S1**). Purified fragments were assembled via CPER, and the CPER product was then directly transfected into BHK/T7-9 cells stably expressing the T7 RNA polymerase^8^ without any DNA purification step. Consistent with the standard RABV RG method, helper plasmids expressing RABV N, P, and L proteins were co-transfected with CPER product to initiate viral genome replication and transcription (**Fig. 1A**). Infectious virus was detected in the culture supernatants from 4 days post-transfection (dpt), with titers increasing up to 10 dpt (**Fig. 1C and 1D**). The culture supernatants harvested at 5 dpt were transferred to neuroblastoma-derived NA cells for further propagation (**Fig. 1A**). Viral genome sequences of the recovered virus were assessed by Sanger sequencing and no nucleotide mutations were observed. CPER-derived RABV exhibited growth kinetics in NA cells comparable to the parental virus (**Fig. 1E**).

**Figure 1.**
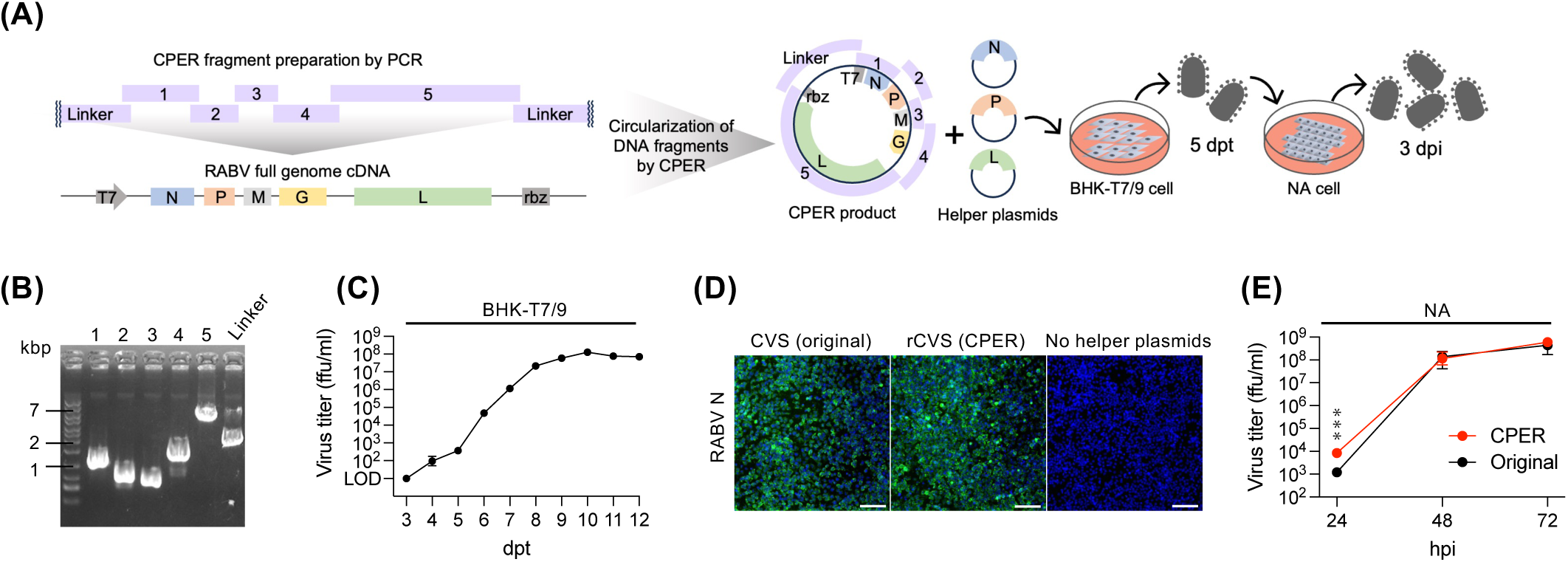
Establishment of a CPER-based reverse genetics system for RABV. (**A**) Schematic image of the CPER design for a wild type RABV and a flow for the recovery of RABV by CPER methods. (**B**) Gel electrophoresis of PCR-amplified CPER fragments. (**C**) Infectious virus titer of RABV in culture medium of BHK/T7-9 cells co-transfected with the CPER product and helper plasmids were measured by focus-forming assay (FFA) at the indicated time points (dpt: days post transfection). (**D**) Detection of RABV N by immunofluorescence assay in NA cells. Culture supernatants from BHK/T7-9 cells transfected with CPER product and helper plasmids were transferred to NA cells at 5 dpt. NA cells were fixed and stained for RABV N at 3 days post infection (dpi). Scale bars: 100 μm. (**E**) Virus growth curves in NA cells. Cells were infected with RABV at a MOI of 0.01, and virus titers in culture media at indicated time points (hpi: hours post infection) were determined by FFA. (**C, E**) Data in the graphs are geometric means ± geometric standard deviations of three replicates from a representative experiment. Statistical analyses: multiple unpaired *t*-tests, \*\*\**P* < 0.001.

### Generation of RABV with a point mutation via CPER

We next tested whether the CPER system could generate RABV carrying a point mutation, using the HEP strain. A mutation substituting glutamine with arginine at position 333 (Q333R) in the glycoprotein (G), which increases virulence of the attenuated HEP strain^25,26^, was introduced by overlap PCR into the G-encoding fragment, which was then included in the CPER assembly (**Fig. 2A**). Both wild-type HEP and HEP with the Q333R mutation (HEP^Q333R^) were successfully rescued (**Fig. 2B**), and Sanger sequencing verified that the rescued HEP^Q333R^ strain retained the intended mutation (**Fig. 2C**). While both viruses showed similar growth kinetics in mouse neuroblastoma-derived NA cells, HEP^Q333R^ strain exhibited impaired growth in human astrocytoma-derived SVG-A cells (**Fig. 2D and 2E**), consistent with our previous observations^25^. These results demonstrate that the CPER system enables efficient generation of biologically relevant RABV mutants.

**Figure 2.**
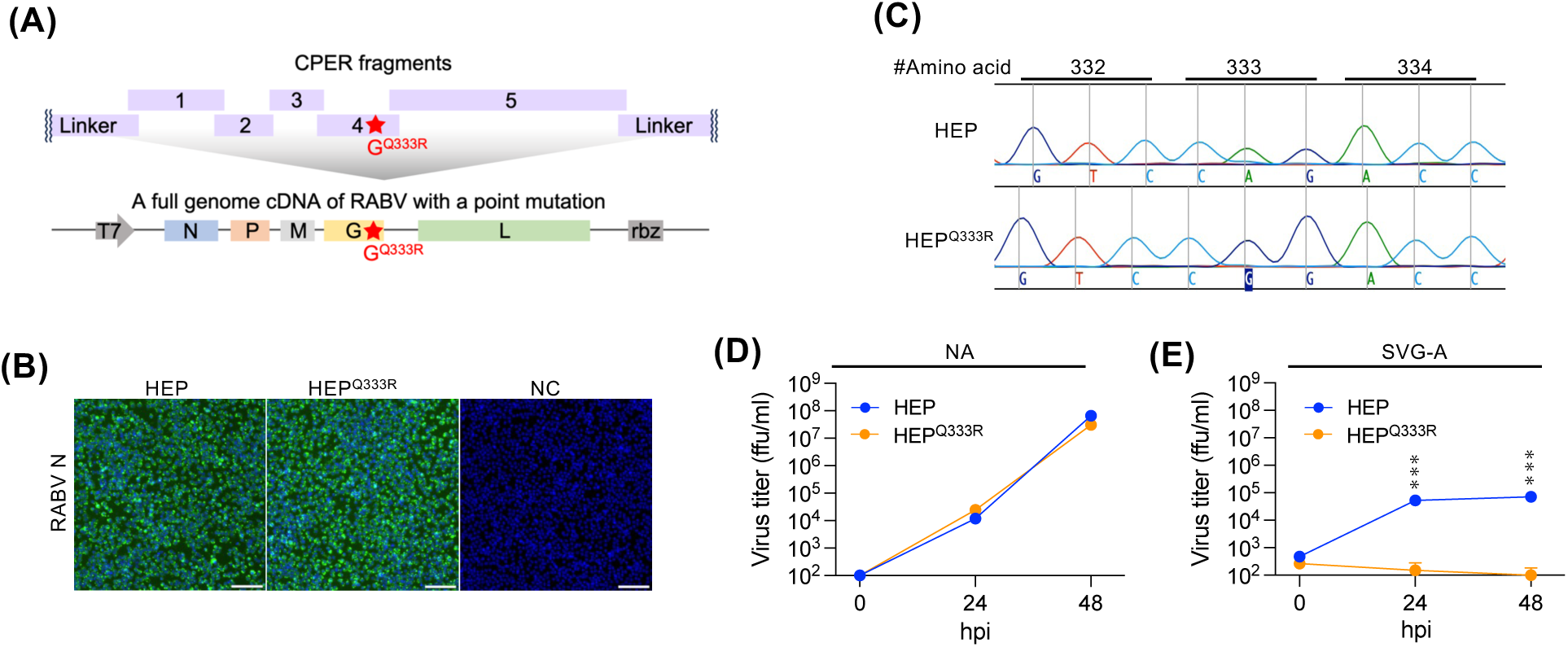
Generation of RABV with a point mutation using CPER. (**A**) Schematic image of the CPER design for RABV harboring the Q333R mutation in the G gene. (**B**) Detection of RABV N by immunofluorescence assay in NA cells. Culture supernatants of the transfected BHK/T7-9 cells were transferred to NA cells at 5 dpt. NA cells were fixed and stained for RABV N at 3 days post infection (dpi). Scale bars: 100 μm. (**C**) DNA sequence electropherograms at amino acid position from 332-334 of the RABV glycoprotein. Viral genomes of CPER-derived RABVs were analyzed by Sanger sequencing. (**D, E**) Virus growth curves in (D) NA and (E) SVG-A cells. Cells were infected with RABV at a MOI of 0.01 for NA and 0.5 for SVG-A cells, and virus titers in the supernatants were measured by FFA at the indicated time points. Data in the graphs are geometric means ± geometric standard deviations of three replicates from a representative experiment. Statistical analysis: multiple unpaired *t*-tests, \*\*\**P* < 0.001.

### Generation of RABV expressing exogenous reporter genes via CPER

One of the advantages of the CPER method is that a foreign gene can be flexibly inserted in the intended position by adding DNA fragments to the reaction. In this study, we designed fragments coding a fluorescent protein GFP or mCherry (NP-mCh) in the intergenic region between the N and P genes, as well as an mCherry fused to the C-terminus of the P gene (P-mCh) (**Fig. 3A**). CPER and transfection were performed in the same manner with the additional reporter gene fragments. After passage in NA cells, the expression of the fluorescent proteins was observed in the cells infected with the recombinant RABVs (**Fig. 3B**). RT-PCR of the viral genome region flanking the insertion site of reporter genes revealed that the reporter genes were stably maintained in the passaged virus populations (**Fig. 3C**). These results validate CPER as a useful method for generating recombinant RABVs expressing exogenous proteins.

**Figure 3.**
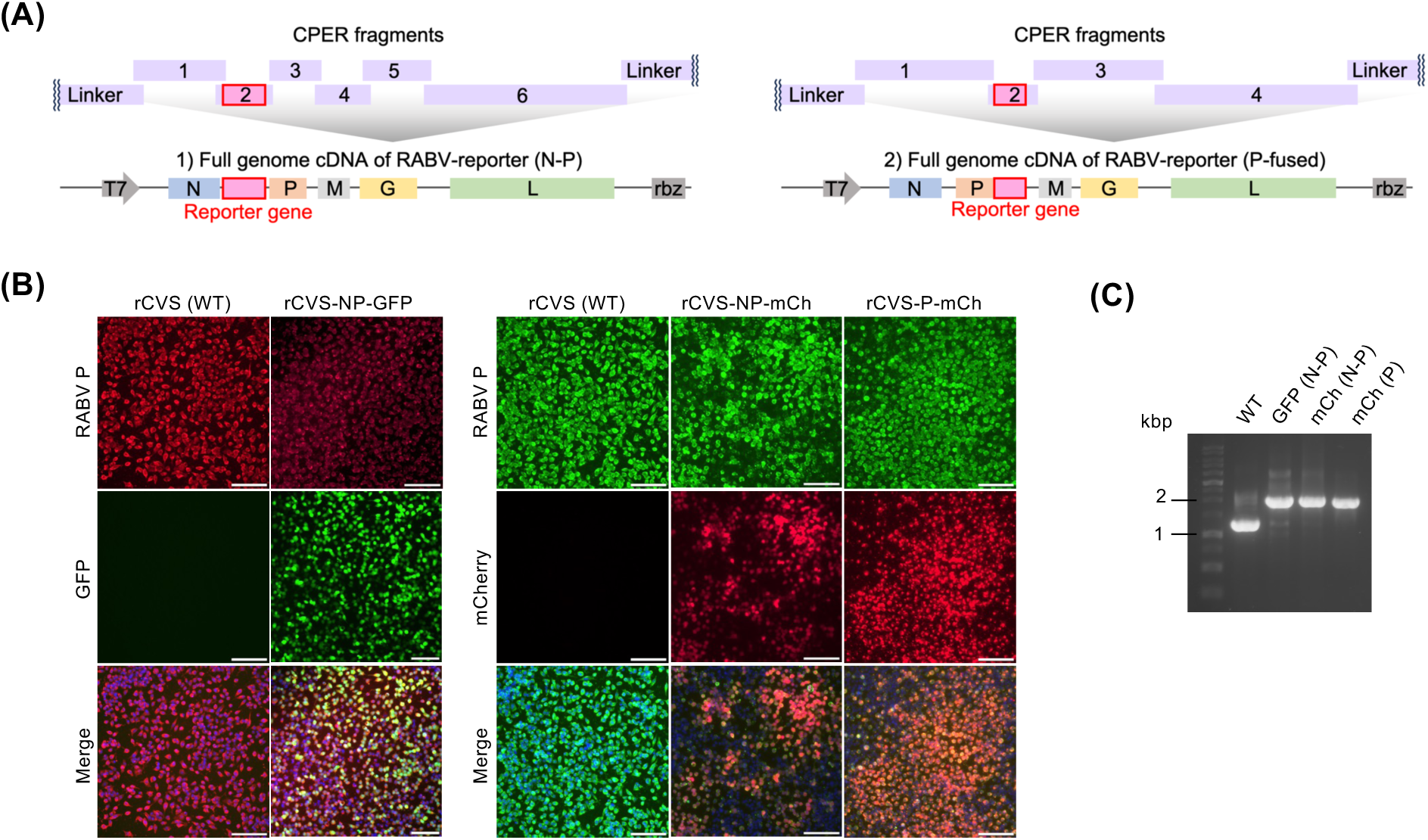
Generation of RABV expressing exogenous reporter genes via CPER. (**A**) Schematic image of the CPER design for RABV with an exogenous reporter gene. (**B**) Detection of RABV P by immunofluorescence assay. NA cells infected with CPER-derived RABVs were fixed and stained for RABV P. Scale bars: 100 μm. (**C**) Gel electrophoresis of RT-PCR amplicons of the viral genome region flanking the insertion site of the reporter genes.

### Generation of chimeric RABVs via CPER

We next generated chimeric RABVs in which the G gene was swapped between the pathogenic CVS and attenuated HEP strains (**Fig. 4A**). The CPER method was successfully applied to recover the chimeric viruses: CVS strain possessing HEP G (CVS-hepG) and HEP strain possessing CVS G (HEP-cvsG) as well as wild-type CVS and HEP strains (**Fig. 4B**). All viruses replicated with similar kinetics in NA cells (**Fig. 4C**). Given the role of RABV G in pathogenicity, we examined virulence of the viruses in a mouse model. Intramuscular inoculation with the highly pathogenic CVS strain resulted in 100% mortality by 5 days post infection (dpi), while all mice inoculated with the attenuated HEP strain survived (**Fig. 4D**). The CVS-hepG chimeric virus produced a delayed disease onset and an increase in survival compared with the parental CVS, whereas HEP-cvsG exhibited enhanced pathogenicity, causing 80% mortality (**Fig. 4D**). These reciprocal phenotypes confirm the critical role of RABV G in strain-specific pathogenicity which is similar to previous studies^27–30^. These results demonstrate how CPER enables facile construction of chimeric viruses for functional analyses.

**Figure 4.**
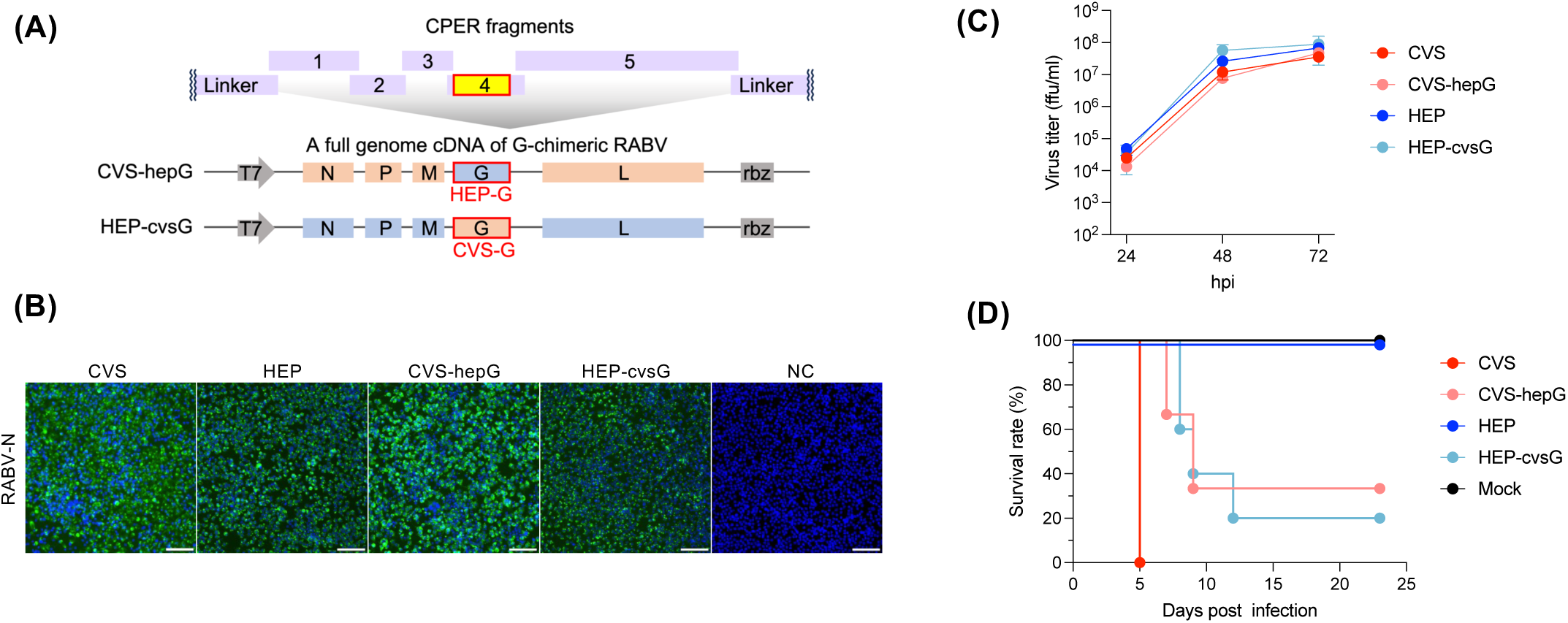
Generation of chimeric RABVs via CPER. (**A**) Schematic image of the CPER design for chimeric RABV. (**B**) Detection of RABV N by immunofluorescence assay. NA cells infected with CPER-derived RABVs were fixed and stained for RABV N. Scale bars: 100 μm. (**C**) Virus growth curves in NA cells. Cells were infected with RABV at a MOI of 0.01, and virus titers in culture media at indicated time points were determined by FFA. Data in the graph are geometric means ± geometric standard deviations of three replicates from a representative experiment. (**D**) Survival curves of mice (n=6) in the RABV infection experiment. Five-week-old ddY mice were intramuscularly inoculated with 5×10^5^ ffu of RABV and monitored daily for survival.

### Deep sequence analysis of CPER-derived RABV and cloning of viral full-genome cDNA via circular polymerase extension cloning (CPEC)

In each experiment, consensus whole genome sequences of CPER-derived RABV genomes were confirmed by Sanger sequencing, as done in other studies^17,21,31^. However, a previous study demonstrated that CPER-derived virus populations could contain low-frequency mutations undetectable by Sanger sequencing analysis^32^. To assess the sequence divergence of the CPER-derived virus populations, we performed deep sequencing analysis on viral RNAs from culture supernatants harvested at 5, 7, 9, and 11 dpt, comparing viruses recovered via conventional plasmid-based and CPER-based RG (n=5 biological replicates each). While plasmid-derived viruses showed no mutations above a 10% frequency, four out of five CPER-derived clones harbored mutations ranging from 10.2% to 57.1% frequency at one or more sites depending on the sampling time points (**Table 1, Table S2**). These mutation patterns were varied, and mutations were not shared among clones (**Table S2**). Since CPER fragments used in this experiment were derived from the same preparation, these results suggest that circularizing PCR can introduce low-frequency mutations, as previously reported for other viruses^32,33^.

**Table 1.** Number of nucleotide substitutions in CPER-derived virus determined by NGS analysis.

|  |  | Frequency of nucleotide substitutions |  |  |  |  |
| --- | --- | --- | --- | --- | --- | --- |
|  |  | 10-20% | 20-30% | 30-40% | 40-50% | >50% |
| CPER1 | 5 dpt | 1 |  |  | 1 |  |
|  | 7 dpt | 5 | 1 | 1 |  |  |
|  | 9 dpt | 2 |  | 2 |  |  |
|  | 11 dpt | 2 |  |  | 2 |  |
| CPER2 | 5 dpt |  |  |  |  |  |
|  | 7 dpt | 1 |  |  |  |  |
|  | 9 dpt |  |  |  |  |  |
|  | 11 dpt |  |  |  |  |  |
| CPER3 | 5 dpt |  |  |  | 2 |  |
|  | 7 dpt |  | 1 |  | 1 |  |
|  | 9 dpt | 1 |  |  |  | 1 |
|  | 11 dpt | 1 |  |  |  | 1 |
| CPER4 | 5 dpt |  |  |  |  |  |
|  | 7 dpt |  |  |  |  |  |
|  | 9 dpt |  |  |  |  |  |
|  | 11 dpt |  |  |  |  |  |
| CPER5 | 5 dpt |  |  |  |  |  |
|  | 7 dpt |  |  |  |  |  |
|  | 9 dpt | 1 |  |  |  |  |
|  | 11 dpt | 3 |  |  |  |  |

\*dpt: days post transfection

To leverage the advantages of CPER and minimize the introduction of unintended mutations during the CPER-based RG, we explored a practical potential to incorporate a cloning step of CPER products to obtain a RABV full-length cDNA plasmid. Purified CPER products with a pUC19 backbone were transformed into *E. coli* HST08 Premium Competent Cells (Takara Bio), and 10 plasmid clones were obtained from individual single colonies. Restriction enzyme digestion demonstrated similar digestion patterns for all clones derived from circular polymerase extension cloning (CPEC) to that of an original plasmid pCVS, indicating that CPER fragments were efficiently circularized to form a circular DNA encoding a full length RABV genome cDNA (**Fig. S1**). Sanger sequencing analysis revealed that 5 out of 10 clones had complete sequences without mutations. A single mutation shared among 4 clones likely originated from the fragment PCR step (**Table 2**). These results indicate that CPEC can serve as a sequence-independent, ligase-free cloning method to generate full-length RABV cDNA plasmids suitable for traditional RG applications.

**Table 2.** Mutations in CPEC-derived plasmids determined by Sanger sequencing.

| Genome region | P |  | G |  | L |  |
| --- | --- | --- | --- | --- | --- | --- |
| Nucleotide position | 1676 | 4638 | 8117 | 8187 | 8869 | 11589 |
| CPEC1 |  |  |  |  |  |  |
| CPEC2 |  |  |  |  | G>A |  |
| CPEC3 | C>T | A>C | C>T |  | G>A |  |
| CPEC4 |  |  |  |  |  |  |
| CPEC5 |  |  |  |  |  | A>G |
| CPEC6 |  |  |  |  | G>A |  |
| CPEC7 |  |  |  |  |  |  |
| CPEC8 |  |  |  |  |  |  |
| CPEC9 |  |  |  | G>A | G>A |  |
| CPEC10 |  |  |  |  |  |  |
| Amino acid change | Q>stop | H>P | S>F | W>stop | D>N | = |

## Discussion

Improvements in RG technologies have significantly advanced virology research. In this study, we established a CPER-based RG platform for RABV. Compared to conventional plasmid-based RG, this approach simplifies and accelerates the generation of recombinant viruses by eliminating the cloning process of a full-length RABV genome cDNA^8,34^. The process from fragment preparation for CPER assembly to transfection of the product can be completed within a single day, enabling more rapid generation of recombinant viruses.

A major advantage of CPER is its flexibility in genetic manipulation^17,19,35^. In our “proof-of-concept” study, we successfully generated recombinant RABVs carrying a point mutation, exogenous reporter genes, and chimeric G genes through simple fragment modification or substitution. CPER strategy avoids sequence constraints that are often required for conventional ligation-based cloning methods and enables flexible and facile construction of recombinant viruses with various genetic modifications. We note that our CPER RG specifically rescued recombinant viruses possessing the intended genetic modifications without contamination of the parental virus.

Importantly our CPER-derived chimeric RABVs possessing a swapped G gene, a well-known virulence determinant, between lethal CVS strain and attenuated HEP strain exhibited incomplete phenotype switching, suggesting that there may be virulence factors for the CVS strain and attenuation factors for HEP strain other than G. Previously, RABV clones with HEP L stably retained G^R333Q^, but those with LEP L underwent a substitution to a high-pathogenic G^Q333R^, indicating HEP L as a possible attenuation factor^36^. A single amino acid substitution of aspartic acid to asparagine at position 80 (D80N) in the matrix protein (M) reportedly increased the virulence of the HEP strain^37^. Considering that pathogenic CVS strain possesses M^D80N^, this might be contributing to the virulence of the CVS strain. Multigenic attenuation properties have been reported based on analysis of chimeric clones between the Nishigahara and RC-HL strains^38^. However, direct comparison of CVS and HEP strains has not been performed in this context. Further studies using chimeric and mutant viruses will help understand viral factors regulating their virulence, where the utility and feasibility of CPER will be highlighted.

Despite the advantages, certain limitations emerged in CPER-driven RG. In this study, both CVS and HEP fixed strains were successfully rescued using CPER; however, the Toyohashi street strain^39^ could not be recovered despite multiple attempts (data not shown). Generally, virus recovery is strongly influenced by the parental strain’s propagation efficiency. However, given that we have successfully rescued the Toyohashi strain using a plasmid-based RG system^34^, the CPER failure of the Toyohashi strain may reflect a lower rescue efficiency, potentially due to suboptimal circularization during assembly. Moreover, deep sequencing revealed that CPER-derived virus populations occasionally contained low-frequency mutations (<50%) undetectable by Sanger sequencing, whereas conventional plasmid-derived viruses did not. This suggests that circularizing PCR in CPER could introduce random mutations into the assembled CPER product due to PCR error, as has also been observed for other CPER-derived viruses^32,33^. Optimization of PCR conditions with high-fidelity DNA polymerase and lower amplification cycles may reduce such experimental artifacts.

The CPER method was originally adapted from CPEC, which bypasses any conventional gene cloning techniques^40^. Here, we further demonstrated that CPER products possessing the replication origin of *E. coli* can be efficiently transformed into *E. coli* to yield full-length RABV cDNA plasmids, offering an alternative route when plasmid-based RG is advantageous for downstream applications.

Until recently, CPER-based RG has been primarily applied to positive-sense RNA viruses, with only a single report describing the recovery of a negative-sense RNA virus, respiratory syncytial virus, by CPER methods^23^. Given the conserved transcription and replication mechanisms across members of the *Mononegavirales*^41^, the CPER-based RG system and its integration with CPEC-based plasmid construction could be broadly applied to other members of the order. This approach has the potential to accelerate the generation and characterization of recombinant mononegaviruses, thereby facilitating both basic virology research and applied studies such as vaccine development and antiviral evaluation.

## Materials and Methods

### Ethics statement

Animal experiments were approved by the Institutional Animal Care and Use Committee of Hokkaido University (approval number 19-0014) and performed according to the committee’s guidelines.

### Cells

Mouse neuroblastoma (NA) cells and baby hamster kidney cells stably expressing T7 RNA polymerase (BHK/T7-9) cells^8^ were maintained in Eagle’s Minimum Essential Medium (MEM) supplemented with 10% fetal bovine serum (FBS). Human fetal astrocyte SVG-A cells were maintained in Dulbecco’s Modified Eagle’s Medium supplemented with 10% FBS. All cells were incubated at 37°C in the presence of 5% CO2.

### Circular polymerase extension reaction (CPER)

DNA fragments were amplified using PrimeSTAR Max (Takara Bio) with primer sets listed in **Table S1**. Fragments were purified via gel extraction using MonoFas (Animos) and equally mixed at 4 nM. Equal volume of PrimeSTAR Max master mix was then added to the fragment mixture. CPER was performed with the following thermal cycling conditions: 95°C for 1 min, 35 cycles of 98°C for 10 s, 71°C for 10 s, 72°C for 80 s, followed by 72°C for 1 min. The CPER products were directly subjected to transfection for virus rescue without any DNA purification steps.

### Virus rescue

BHK/T7-9 cells ^8^seeded in a 12-well plate were transfected with either 4 μg of a plasmid encoding full genome cDNA of RABV (pCVS)^24^ or 25 μl of unpurified CPER product together with helper plasmids: 0.8 μg of pT7/IRES-RN, 0.2 μg of pT7/IRES-RP, and 0.4 μg of pT7/IRES-RL^8^ using 16.2 μl of TransIT-LT1 Transfection Reagent (Mirus Bio). Five days after transfection, culture supernatants were transferred to NA cells, and virus were collected at 3 days later. All viruses were propagated in NA cells and virus stocks were stored at −80℃.

### Immunofluorescence staining

Cells were fixed with formalin at room temperature overnight and washed with PBS. Fixed cells were incubated at room temperature for 1 hour in 0.4% Block ACE (KAC) 0.1% Tween-20 in PBS with the following antibodies: FITC-RABV N (1:400, #800-092, Fujirebio), RABV P (1:1,000, #A54523-100, EpiGentek). After washing three times with 0.01% Tween 20 in PBS, the following secondary antibodies were used for indirect fluorescence assays under the same conditions as the first antibody; Alexa Fluor 488-anti-mouse IgG (1:1,000, #A-11001, Invitrogen), Alexa Fluor 594-anti-rabbit IgG (1:1,000, #A-11012, Invitrogen). For nuclear staining, Hoechest 33342 (5 μg/ml) was added to the staining solution.

### Deep sequencing analysis

After introducing CPER products and helper plasmids into BHK/T7-9 cells, the cells were washed with PBS and refed with fresh medium on the following day to completely remove the input transfection complex. Cell culture supernatants harvested at 5, 7, 9, and 11 dpt were subjected to RNA extraction using TRIzol LS (Invitrogen) and Direct-zol RNA Miniprep kit (Zymo Research). The RABV full genome was amplified by RT-PCR into three segments with primer sets listed in **Table S3** using PrimeScript One Step RT-PCR Kit Ver.2 (Takara Bio) with the following conditions: 50℃ for 30 min, 94℃ for 2 min, and 40 cycles of 94℃ for 30 sec, 68℃ for 30 sec, 72℃ for 3 min 30 sec. PCR amplicons were purified by using KAPA HyperPure beads (Roche). Purified amplicons originating from the same RNA samples were pooled at equal concentrations. DNA libraries were constructed using Illumina DNA PCR-Free Prep (Illumina), and 300-bp paired-end sequencing was performed on Illumina MiSeq (Illumina). Sequence reads were trimmed and assembled on a reference RABV sequence (GenBank accession No. LC325820.1), followed by mutation analysis to extract mutations with >10% frequency using the CLC Genomics Workbench 23 (Qiagen).

### Virus titration

Virus titers were determined by the focus-forming assay (FFA). Serial-diluted samples were inoculated into NA cells seeded in 48-well plates. After 1 h-incubation, samples were removed and cells were overlayed with MEM supplemented with 5% FBS, 0.5% methyl cellulose, and GlutaMAX (Gibco). Following 3 days of incubation, the cells were fixed and stained with FITC-labelled anti-RABV N antibody (1:300, #800-092, Fujirebio) and Hoechst 33342 (5 μg/ml). Foci were counted, and the virus titers were described as focus-forming units (ffu).

### Circular polymerase extension cloning (CPEC)

Purified CPER products with KAPA HyperPure beads (Roche) were transformed into the *E. coli* HST08 Premium Competent Cells (Takara Bio), and *E. coli* was cultured on Luria-Bertani (LB) agar plates with 100 μg/ml ampicillin at 37℃ for 16 hours. Single colonies were picked for LB broth liquid culture at 37℃ for 16 hours, and plasmids were extracted using the QIAprep Spin Miniprep Kit (QIAGEN).

### Viral growth curves

NA cells and SVG-A cells were infected with viruses at a multiplicity of infection (MOI) of 0.01 and 0.5, respectively. Supernatants were collected at the indicated time points in the Figures, and virus titers were measured by FFA.

### Animal experiments

Five-week-old male ddY mice were intramuscularly inoculated under anesthesia using isoflurane. Animals were observed daily for disease symptoms until they reached the humane endpoint defined as a 20% decrease in body weight or an inability to reach food or water due to the onset of disease.

### Statistical analysis

Statistical analyses were performed using GraphPad Prism 10.1.1. Multiple unpaired *t*-tests with the Benjamini, Krieger, and Yekutieli method were performed for comparisons of two groups at multiple time points. Data are presented as geometric means ± geometric standard deviations in graphs. \*\*\**P* < 0.001.

### Data availability

This study includes no data deposited in external repositories.

## Acknowledgments

This study was supported in part by the Japan Society for the Promotion of Science (JSPS) KAKENHI under grant numbers 23H02376, 23K19326, 18KK0192; and the Japan Agency for Medical Research and Development (AMED) under grant numbers JP223fa627005, JP24wm0225044, JP20wm0125008.

## Author Contributions

Y.I. and M.S. designed the research; Y.I., N.K., K.T., G.G., K.K., A.O., I.F. and M.S. performed experiments and analyzed data; N.I., S.S., W.H., Y.O. and H.S. provided critical reagents and advice; Y.I. and M.S. wrote the draft manuscript; N.I., S.S., W.H., Y.O. and H.S. provided critical reviews of the draft manuscript; Y.I., N.I., H.S. and M.S. acquired funding supports.

## Figure Legends

**Figure S1.** Evaluation of CPEC-derived plasmids coding a full-genome cDNA of RABV. Gel electrophoresis of CPEC-derived plasmids digested by the restriction enzyme *EcoRI*. Ct: control parental plasmid pCVS.

