## Supplemental files for "Application of the CPER reverse genetics system for genetic engineering of rabies virus"

### Figure S1

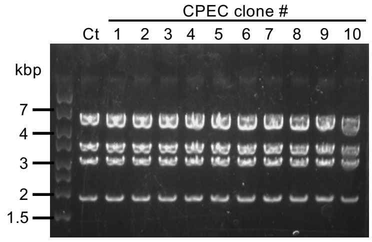


**Figure S1. Evaluation of CPEC-derived plasmids coding a full-genome cDNA of RABV.**

Gel electrophoresis of CPEC-derived plasmids digested by a restriction enzyme *EcoRI*. Ct: control parental plasmid pCVS.

### Table S1

**Table S1**. Primers used for the preparation of CPER DNA fragments in this study

| Target virus | Fragment | Primer name | Type[a] | Sequence (5'-3') |
|---|---|---|---|---|
| CVS | CVS-N | CVS-CP-N-F | F | TAATACGACTCACTATAGGGACGCTTAACAACAAAACCAGAGAAGAAAAAGACAG |
| | | CVS-CP-N-R | R | CATATTTGGGATGGTTTGAAAGGAGGAGTGTTAGTTTTTTTCATGATGGATATATAC |
| | CVS-P | CVS-CP-P-F | F | GTATATATCCATCATGAAAAAAACTAACACTCCTCCTTTCAAACCATCCCAAATATG |
| | | CVS-CP-P-R | R | AACGTTCATTTTATCAGTGGTGTTGCCTGTTTTTTTCATGTCTACTCCATAAC |
| | CVS-M | CVS-CP-M-F | F | GTTATGGAGTAGACATGAAAAAAACAGGCAACACCACTGATAAAATGAACGTT |
| | | CVS-CP-M-R | R | CCTTAAGTCTTTTGAGGGATGTTAATAGTTTTTTTCACATCCAAGAGGCTCAA |
| | CVS-G | CVS-CP-G-F | F | TTGAGCCTCTTGGATGTGAAAAAAACTATTAACATCCCTCAAAAGACTTAAGG |
| | | CVS-CP-G-R | R | GTTCTTTTGGTCTACAGTTTTTTTCTCGACTGAAATGCTTAG |
| | CVS-L | CVS-CP-L-F | F | CTAAGCATTTCAGTCGAGAAAAAAACTGTAGACCAAAAGAAC |
| | | CVS-CP-L-R | R | GTCCGGATTCAAGATCTTGTTTTTTTCAAGATGCATCATACAAGA |
| | CVS-Linker | CVS-CP_Linker-F | F | TCTTGTATGATGCATCTTGAAAAAAACAAGATCTTGAATCCGGAC |
| | | CVS-CP_Linker-R | R | CTGTCTTTTTCTTCTCTGGTTTTGTTGTTAAGCGTCCCTATAGTGAGTCGTATTA |
| CVS-mCherry (N-P) | CVS-N-mCherry | CVS-CP-N-F | F | TAATACGACTCACTATAGGGACGCTTAACAACAAAACCAGAGAAGAAAAAGACAG |
| | | CVS_CPER1-R | R | GTTTGAAAGGAGGAGTGTTAGTTTTTTTCATGATGGATATATACAATCT |
| | CVS-mCherry | CVS_NP_mCherry-F | F | TGTATATATCCATCATGAAAAAAACTAACACTCCTCCTTTCAAACCATCCCAAATCGCCACCATGGTGAGCAAGGGCGAGGAG |
| | | CVS_NP_mCherry-R | R | ATTTGGGATGGTTTGAAAGGAGGAGTGTTAGTTTTTTTCACTTGTACAGCTCGTCCATGCCGCCGGTG |
| | mCherry-CVS-P/M/G | CVS_CPER2-F | F | CAAACCATCCCAAATATGAGCAAGATCTTTGTTA |
| | | CVS-CP-G-R | R | GTTCTTTTGGTCTACAGTTTTTTTCTCGACTGAAATGCTTAG |
| CVS-mCherry (P-fused) | CVS-P-mCherry | CVS-CP-N-F | F | TAATACGACTCACTATAGGGACGCTTAACAACAAAACCAGAGAAGAAAAAGACAG |
| | | CVS_CPER_m1-R | R | TTATCCTCCTCGCCCTTGCTCACCATGCAGGATGTATAGCGATTCAAATCATCTTG |
| | CVS-Pfusion-mCherry | CVS_P_mCherry-F | F | TGAATCGCTATACATCCTGCATGGTGAGCAAGGGCGAGGAGGATAACATGG |
| | | CVS_P_mCherry-R | R | GGACTGAGTTCGAAAACTCGGTTACTTGTACAGCTCGTCCATGCCGCCGGTGGAGTG |
| | CVS-mCherry-M/G | CVS_CPER_m2-F | F | GGCATGGACGAGCTGTACAAGTAACCGAGTTTTCGAACTCAGTCCCTCCAGATAATGA |
| | | CVS-CP-G-R | R | GTTCTTTTGGTCTACAGTTTTTTTCTCGACTGAAATGCTTAG |
| CVS-GFP (N-P) | CVS-N-GFP | CVS-CP-N-F | F | TAATACGACTCACTATAGGGACGCTTAACAACAAAACCAGAGAAGAAAAAGACAG |
| | | CVS_CPER1-R | R | GTTTGAAAGGAGGAGTGTTAGTTTTTTTCATGATGGATATATACAATCT |
| | CVS-GFP | CVS_NP_AcGFP-F | F | TGTATATATCCATCATGAAAAAAACTAACACTCCTCCTTTCAAACCATCCCAAATCGCCACCATGGTGAGCAAGGGCGCCGAGCTGT |
| | | CVS_NP_AcGFP-R | R | TAACAAAGATCTTGCTCATATTTGGGATGGTTTGAAAGGAGGAGTGTTAGTTTTTTTCATTACTTGTACAGCTCATCCATGCCGTG |
| | GFP-CVS-P | CVS_CPER2-F | F | CAAACCATCCCAAATATGAGCAAGATCTTTGTTA |
| | | CVS-CP-P-R | R | AACGTTCATTTTATCAGTGGTGTTGCCTGTTTTTTTCATGTCTACTCCATAAC |
| HEP | HEP-N | HEP-CP-N-F | F | ACGCTTAACAACAAAACCAAAGAAGAAGCAGACATCGTCAGTTGCAAAGCAAAAA |
| | | HEP-CP-N-R | R | ACTTGGGATGGTTCGAAAGGAGGAGTGTTAGTTTTTTTCA |
| | HEP-P | HEP-CP-P-F | F | TGAAAAAAACTAACACTCCTCCTTTCGAACCATCCCAAGT |
| | | HEP-CP-P-R | R | GTTCATTTTATTAGTGGTGTTGCCTGTTTTTTTTCATATCGACTCCAT |
| | HEP-M | HEP-CP-M-F | F | ATGGAGTCGATATGAAAAAAAACAGGCAACACCACTAATAAAATGAAC |
| | | HEP-CP-M-R | R | AGGAACCATCTTTCCTTAAGTCTTTTGAGGGATGTTAATAGTTTTTTTCACAT |
| | HEP-G | HEP-CP-G-F | F | ATGTGAAAAAAACTATTAACATCCCTCAAAAGACTTAAGGAAAGATGGTTCCT |
| | | HEP-CP-G-R | R | TCCCGGATCCAGCATCTTGATATGGGTCTCGAGATGAGAAGT |
| | HEP-L | HEP-CP-L-F | F | ACTTCTCATCTCGAGACCCATATCAAGATGCTGGATCCGGGA |
| | | HEP-CP-L-R | R | GATTCAAGATCTTGTTTTTTTCAAGATGCATCATACAAGA |
| | HEP-Linker | HEP-CP_Linker-F | F | TCTTGTATGATGCATCTTGAAAAAAACAAGATCTTGAATC |
| | | HEP-CP_Linker-R | R | TTTTTGCTTTGCAACTGACGATGTCTGCTTCTTCTTTGGTTTTGTTGTTAAGCGT |
| CVS-hepG | CVS-hepG-M | CVS-CP-M-F | F | GTTATGGAGTAGACATGAAAAAAACAGGCAACACCACTGATAAAATGAACGTT |
| | | cvsM-R-hepG | R | CCTTAAGTCTTTTGAGGGATGTTAATAGTTTTTTTCACATCCAAGAGGCTCAA |
| | CVS-hepG-G | cvsM-F-hepG | F | TTGAGCCTCTTGGATGTGAAAAAAACTATTAACATCCCTCAAAAGACTTAAGG |
| | | hepG-R-cvsL | R | GTTCTTTTGGTCTACAGTTTTTTTCTCGACTGAAATGCTTAG |
| | CVS-hepG-L | hepG-F-cvsL | F | CTAAGCATTTCAGTCGAGAAAAAAACTGTAGACCAAAAGAAC |
| | | CVS-CP-L-R | R | GTCCGGATTCAAGATCTTGTTTTTTTCAAGATGCATCATACAAGA |
| HEP-cvsG | HEP-cvsG-M | HEP-CP-M-F | F | ATGGAGTCGATATGAAAAAAAACAGGCAACACCACTAATAAAATGAAC |
| | | hepM-R-cvsG | R | CCTTAAGTCTTTTGAGGGATGTTAATAGTTTTTTTCACATCCAAGAGGCTCAA |
| | HEP-cvsG-G | hepM-F-cvsG | F | TTGAGCCTCTTGGATGTGAAAAAAACTATTAACATCCCTCAAAAGACTTAAGG |
| | | cvsG-R-hepL | R | GTTCTTTTGGTATACAGTTTTTTTCTCGACTGAAATGCTTAG |
| | HEP-cvsG-L | cvsG-F-hepL | F | CTAAGCATTTCAGTCGAGAAAAAAACTGTATACCAAAAGAAC |
| | | HEP-CP-L-R | R | GATTCAAGATCTTGTTTTTTTCAAGATGCATCATACAAGA |

[a]F; forward primer, R; reverse primer

### Table S2

**Table S2**. Mutations and frequency (%) in CPER-derived virus determined by NGS analysis

| Genome region | | N | | | IGR | P | IGR | G | | L | | | | |
|---|---|---|---|---|---|---|---|---|---|---|---|---|---|---|
| Nucleotide position | | 139 | 1182 | 1247 | 1484 | 1919 | 2410 | 3824 | 4750 | 6020 | 6333 | 8995 | 9616 | 10293 |
| Reference nucleotide | | G | C | G | T | C | G | C | G | C | C | T | G | C |
| Nucleotide change | | A | T | T | C | A | T | T | T | T | A | C | T | T |
| Amino acid change | | = | A>V | D>Y | N/A | P>T | N/A | H>Y | M>I | T>I | N>K | = | G>W | = |
| CPER1 | 5 dpt | | | | | | | 47.1 | | | | | 17.4 | |
| | 7 dpt | | | 29.3 | 10.4 | | | 14.4 | 13.5 | 10.5 | | | 31.2 | 11.3 |
| | 9 dpt | | | 39.4 | | | | | 12.8 | 11.3 | | | 36.7 | |
| | 11 dpt | | | 47.2 | | | | 10.2 | 11.2 | | | | 45.4 | |
| CPER2 | 5 dpt | | | | | | | | | | | | | |
| | 7 dpt | | 11.4 | | | | | | | | | | | |
| | 9 dpt | | | | | | | | | | | | | |
| | 11 dpt | | | | | | | | | | | | | |
| CPER3 | 5 dpt | 40.6 | | | | 46.8 | | | | | | | | |
| | 7 dpt | 26.7 | | | | 47.3 | | | | | | | | |
| | 9 dpt | 16.9 | | | | 55.5 | | | | | | | | |
| | 11 dpt | 13.6 | | | | 57.1 | | | | | | | | |
| CPER4 | 5 dpt | | | | | | | | | | | | | |
| | 7 dpt | | | | | | | | | | | | | |
| | 9 dpt | | | | | | | | | | | | | |
| | 11 dpt | | | | | | | | | | | | | |
| CPER5 | 5 dpt | | | | | | | | | | | | | |
| | 7 dpt | | | | | | | | | | | | | |
| | 9 dpt | | | | | | 11.4 | | | | | | | |
| | 11 dpt | | | | | | 10.4 | | | | 12.8 | 10.8 | | |

### Table S3

**Table S3**. Primers used for library preparation for the NGS analysis

| Primers | Sense | Sequence (5' - 3') | Anealing site positions [a] |
|---|---|---|---|
| CVS-frag1(+) | + | ATACGACTCACTATAGGGACGCTTAACAACAAAACCAGAG | 1-22 |
| CVS-frag1(-) | - | GTCATATGGGTCCAAATCTGTCACACTTGGGGATATGATA | 3814-3775 |
| CVS-frag2(+) | + | GGTCTGAAGAGGACAAAGACTCTTCTCTGCTTCTAGAATA | 3064-3103 |
| CVS-frag2(-) | - | CATTATATTGGCGAGGTTGACTATTTGGTCGTTAGAGATG | 7893-7854 |
| CVS-frag3(+) | + | AGAGTTTTTGAAATCTATAGACCTCGGAGGATTGCCAGAT | 6972-7011 |
| CVS-frag3(-) | - | CAAGATGCATCATACAAGAATTTTAGCATGCACAGGCTTT | 11850-11811 |

a: Based on the genome nucleotide number of rabies virus CVS strain (GenBank accession No.LC325820.1).
